# *Drosophila* dTBCE recruits tubulin around chromatin to promote mitotic spindle assembly

**DOI:** 10.1101/2020.01.21.912428

**Authors:** Mathieu Métivier, Emmanuel Gallaud, Aude Pascal, Jean-Philippe Gagné, Guy G. Poirier, Denis Chrétien, Romain Gibeaux, Laurent Richard-Parpaillon, Christelle Benaud, Régis Giet

## Abstract

Proper assembly of mitotic spindles requires microtubule nucleation at centrosomes but also around chromatin. In this study, we reveal a novel mechanism by which an enrichment of tubulin in the nuclear space following nuclear envelope breakdown promotes nucleation of spindle microtubules. This event mediated by the tubulin-specific chaperone dTBCE, depends on its tubulin binding CAP-Gly motif and is regulated by Ran. Live imaging, proteomic and biochemical analyses suggest that dTBCE is enriched in the nucleus at nuclear envelope breakdown and interacts with nuclear pore proteins and the Ran machinery to create an environment that facilitates subsequent tubulin enrichment. We propose that dTBCE-dependent increase in tubulin concentration in the nuclear space is an important mechanism for microtubule nucleation in organisms where compartmentalization prevents free diffusion of tubulin.

## Introduction

Microtubules (MTs) are essential dynamic components of the cytoskeleton that assemble from α− and β-tubulin dimers ^1^. During cell division, the MT cytoskeleton is remodeled to assemble a MT-based bipolar spindle that ensures the accurate attachment of sister chromatids to the opposite poles and allow their equal segregation toward the two daughter cells. *In vitro*, nucleation of MTs happens spontaneously in the presence of GTP when the concentration of tubulin dimers reaches a critical concentration. In somatic cells, the nucleation of spindle MTs is more complex and occurs both at the centrosomes and around mitotic chromatin during mitosis. MT nucleation during spindle assembly requires several additional factors. These MT nucleating protein complexes include the gamma tubulin ring small complex (γTuSC), which is targeted to the centrosome and preexisting MTs ^2, 3, 4, 5^. MT nucleation is also triggered around chromatin by a Ran-GTP gradient that allows the local release of Spindle Assembly Factors (SAFs) sequestered by importins ^6^. However, the Ran pathway is not essential for MT nucleation in some model organisms ^7, 8^. Indeed, the Chromosome Passenger Complex (CPC) also contributes to MT nucleation and can compensate the absence of Ran ^9, 10^. Interestingly, even if *in vitro* MT nucleation is known to be dependent upon tubulin concentration, local accumulation of free tubulin has only recently started to be considered as a key element that could significantly contribute to spindle assembly ^11, 12^. Surprisingly, in *C.elegans* and *D. melanogaster* an accumulation of free tubulin around centrosomes and chromosomes has been observed during mitosis, but the physiological relevance of this local tubulin increase remains elusive ^11, 13–15^.

Tubulin-binding co-factors (TBCs) exist as five proteins termed TBCA to TBCE, whose concerted action leads to the assembly of polymerization-competent tubulin heterodimers ^16^. However, whether TBCs have independent functions remains an open question. Interestingly, we have previously shown that the knock down of *Drosophila melanogaster* tubulin-binding co-factor E (dTBCE) specifically produced abnormal mitoses in brain neuroblasts (Nbs) ^17^. Here, we demonstrate that dTBCE is responsible during mitosis for the recruitment of tubulin around chromatin following NEBD (nuclear envelope breakdown), and this mechanism is essential for proper spindle assembly. This process represents a novel and likely conserved mechanism that may serve to control MT assembly in space and time.

## Material and methods

### DNA constructs

dTBCE open reading frame was amplified by PCR using the GM13256 cDNA gold clone obtained from the Drosophila Genomic Resource Center (DGRC, Indiana University) and cloned without the stop codon into pENTR (Invitrogen) to generate the pENTR-dTBCE entry clone. This clone was subsequently used for mutagenesis using the QuickChange Ligthning site-directed mutagenesis kit (Agilent). The putative weak NLS motif GLVM<u>KRFR</u>LS detected in the C-terminal domain of dTBCE was mutated into GLVM<u>AAFA</u>LS to generate pENTR-dTBCE^mutNLS^. The HsTBCE deletion nearby the CAP-Gly domain, observed in patients exhibiting hypoparathyroidism, mental retardation and facial dysmorphism (HMF) was recapitulated by deleting the corresponding IVDG aminoacids sequence of dTBCE to generate pENTR-dTBCE^HMF 18^. The conserved CAP-Gly GLRG<u>KH</u>NG motif present in the N-terminal region of dTBCE sequence was mutated into GLRG<u>AA</u>NG (Michel Steinmetz, personal communication) to generate pENTR-dTBCE^mutCG^. To generate C-terminal GFP fusion of wild type and mutant dTBCE proteins, the corresponding mutated entry clones were recombined using a LR clonase kit (Invitrogen) into pUWG (for dTBCE), pTWV (for dTBCE, dTBCE^HMF^, dTBCE^mutCG^ and dTBCE^mutNLS^), or pPWG (dTBCE, dTBCE^HMF^, dTBCE^mutCG^ and dTBCE^mutNLS^) obtained from the DGRC. The destination vectors were subsequently injected into *Drosophila* embryos for P-element mediated transformation (BestGene). Note that the pUWG destination vector allowed the expression of wild type dTBCE under the control of the poly-ubiquitin promoter, while pPWG/V and pTWG/V were used for overexpression with the Gal4/UAS system.

The N-terminal domain of dTBCE (amino acid 1 to 207) was cloned using the Infusion system (Takara) into pET23B (Clontech) to generate the pET23b-Nt-dTBCE vector for expression in *E. coli* and antibody production in guinea pig.

For generation of S2 stable cell lines expressing dTBCE-GFP or dTBCE^mutCG^-GFP, the corresponding entry clones were recombined into pPURO-GFP, allowing expression of C-terminal GFP fusion proteins expressed under the control of the actin 5C promoter, and selection by puromycin (kind gift of Roger Karess, Institut Jaques Monod, Paris).

### Fly stocks

Flies were maintained under standard conditions at 25°C. The *dtbce* mutants flies *tbce^LH15^* and *tbce^zz0241^* have been described previously and correspond to null alleles with embryonic lethality ^19^. The *tbce^LH15^*/*tbce^ZZ0241^* heteroallelic combination was used in this paper and referred as *dtbce* to assay for rescue by the GFP-tagged dTBCE variants.

Expression of dTBCE-GFP under the control of a polyubiquitin promoter was able to rescue both the viability and the fertility of the *dtbce* mutant. Transgenic flies with the following genotypes were used for RNAi or protein expression: TubGal80^ts^; 69B-Gal4, Insc-Gal4, UASt-Cherry-α-tubulin, actin5C-Gal4, UAS-CD8-GFP and V32C-Gal4 and were obtained from the Bloomington Drosophila Stock Center (Indiana University). Nup107-RFP and lamin-GFP lines were described before ^22, 23^. β-tubulin-GFP flies were described before ^20^. UASt or UASp mediated expression of dTBCE-GFP, dTBCE^mutCG^-GFP, dTBCE^mutNLS^-GFP or dTBCE^HRD^-GFP was driven by 69B-Gal4, Actin5C-Gal4 or V32C-Gal4. Transgenic flies carrying UAS and hairpin sequences to knock down dTBCE (Line KK105246 and GD 34388) and Ran (Line GD24835), were obtained from the Vienna *Drosophila* RNAi Center (VDRC) ^21^. Flies expressing the full-length EB1-GFP were described in a previous study ^17^. dTBCE-GFP, dTBCE^mutNLS^-GFP or dTBCE^HMR^-GFP expression (in pUASt and pUASp) driven by the Actin5C driver were able to rescue the lethality of the *dtbce* mutation, by contrast to the expression of dTBCE^mutCG^-GFP.

### Expression and purification of recombinant protein

The pET23b-Nt-dTBCE expression vector was transformed into the *E. coli* BL21(DE3) strain (New England Biolabs). The recombinant protein expression was induced with 1mM IPTG in cells during 4 h at 25°C and the bacterial pellet was stored at −20°C until use. The pellet was resuspended on ice, in Lysis Buffer (LB: Tris 10 mM, 10 % Glycerol, 500 mM NaCl, pH7.4) containing 1% Triton X-100 (Sigma), proteases inhibitor tablets (Roche) and lysozyme (1mg/ml). The resuspended bacterial pellet was then sonicated 10 times for 10 s on ice and centrifuged at 10 000 *g* for 30 min at 4°C. The 6xHistidine-tagged Nt-dTBCE protein in the supernatant was purified following the manufacturer’s instructions on a Nickel column (Quiagen), dialyzed overnight in PBS at 4°C and used for guinea pig immunization (Covalab).

### Antibodies

Polyclonal rabbit anti-Ran antibody (ref. ab11693) was from Abcam and used 1:2000 for Western blotting. The YL1/2 rat monoclonal anti-tyrosylated tubulin antibody (ref Mab1864, 1:200) and the mouse monoclonal and rabbit polyclonal anti-phosphorylated histone H3 (Ser10) antibodies (ref. 06570, 1:1000) were obtained from Millipore. The anti-GFP mouse monoclonal antibody mix (ref. 11814460001, 1:1000) was obtained from Roche. The rabbit anti-PKCζ (ref. C-20, 1:200), the mouse anti-β tubulin antibody (ref. sc-365791, 1:1000) and rabbit anti-actin (ref. sc-1616, 1:5000) polyclonal antibodies were obtained from Santa Cruz. Mouse anti-lamin (ref. ADL04, 1:100) was from purchased from the Drosophila Studies Hybridoma Bank. The mouse monoclonal anti-dTBCE antibody was described before ^19^. The guinea pig polyclonal anti-Nt-dTBCE (Amino acids 1-207) was used at 1:1000 for immunostaining and at 1:10000 for Western blotting. The mouse DM1α monoclonal anti-α-tubulin antibody was from Sigma (ref. T6199, 1:2000). Goat secondary peroxidase-conjugated antibodies (1:5000) were obtained from Jackson ImmunoResearch Laboratories, and donkey Alexa Fluor–conjugated secondary antibodies (1:1000) were obtained from Life Technologies.

### Western Blotting analyses

After transfer of the proteins on a nitrocellulose membrane (GE Healthcare) using a Trans Blot Turbo system (Biorad), the membranes were briefly stained with ponceau S, to ensure sample quality and blocked for 2 h with TBST (20 mM Tris, 150 mM NaCl and 0.1 % Tween20, pH 7.4) containing 10 % skimmed milk (or 5% BSA for the anti-Ran antibody). Primary antibody incubation was performed overnight at 4°C in TBST containing 5 % skimmed milk (or 2.5% BSA for the anti-Ran antibody). Incubation with the secondary antibodies was performed in 5 % skimmed milk for 2 h at room temperature. Three 15 min washes in TBST were performed following each antibody incubation. For western blotting, ECL reagent (ref. 34095) was purchased from Thermo Fisher Scientific.

### Immunofluorescence analyses

Immunostaining of brain neuroblasts (Nbs), MT recovery after cold treatment and image acquisition with a SP5 confocal microscope (Leica) were performed as described previously ^17^.

### Live microscopy

Fresh 0-2 h embryos expressing fluorescent proteins were immobilized on double-sided tape and gently dechorionated by hand with fine point forceps. Two to three embryos were then deposited at the top a drop of halocarbon 700 oil (Sigma) on a glass slide separated by two 22×22 mm coverslips to create a spacer. The embryos were then covered by a 40×22 mm coverslip right before imaging.

Brains expressing the various fluorescent proteins were dissected in Schneider’s *Drosophila* medium containing 10% FCS, following a protocol described before ^17^. To monitor tubulin import in the nuclear space the MT network was depolymerized by 10 min incubation in 20 μM colchicine (Sigma). The preparations were then sealed with mineral oil (ref. M8410, Sigma). Alternatively, brains were dissected in PBS and loaded on a glass slide. The tissue was then gently flattened in 10 μl of PBS with a 22×22 mm coverslip, the excess of buffer was removed and the preparation was sealed by halocarbon 700 oil to avoid evaporation. The Nbs at the border of the tissue were identified by light microscopy and immediately imaged within 15 min. For bead fluorescence analyses, GFP, dTBCE-GFP and dTBCE^mutCG^-GFP beads were incubated in a 20 μM solution of bovine brain tubulin (PurSolutions, ref: 142001) supplemented with 25 % of tubulin-rhodamin (Tebu) and 1 mM GTP in BRB80 (80mM PIPES, 1mM EGTA, 1mM MgCl_2_, pH 6.8 with KOH).

Live images were acquired with a spinning-disk system mounted on an inverted microscope (Elipse Ti; Nikon) equipped with an ILAS2 FRAP head and a 60X 1.4 NA objective at 25°C. Z series were acquired every 1, 30 or 60 s with a CCD camera (CoolSnap HQ2; Photometrics) and a sCMOS ORCA Flash 4.0 (Hamamatsu) controlled by MetaMorph acquisition software version X. Images were processed with ImageJ software ^25^.

### Immunoprecipitation

Stable S2 cells expressing GFP, dTBCE-GFP and dTBCE^mutCG^-GFP were grown and collected by centrifugation at 500 *g* for 2 min. The cells were gently resuspended in PBS at 500 *g* for 2 min, the cell pellets were flash frozen in liquid nitrogen and stored at −80°C or processed immediately for immunoprecipitation. A cell pellet, corresponding to 10^8^ cells was resuspended in 0.5 ml of lysis buffer containing 10 mM Tris, 150 mM NaCl, 0.5 mM EDTA, protease and phosphatase inhibitors (Roche). The resuspended cell pellet was sonicated 10 times for 10 s on ice and centrifuged at 10 000 *g* for 30 min at 4°C. The supernatant was then incubated with 10 μl of a GFP-TRAP-MA beads that were previously washed with lysis buffer (Chromotek). Immunoprecipitation was performed for 1 h at 4°C. The beads were then subjected to 3 washes for 5 min each, transferred in a new tube and analyzed by Western blotting or subjected to proteomic analyses. For tubulin *in vitro* binding assays, we collected *Drosophila* embryos from females expressing GFP, dTBCE-GFP and dTBCE^mutCG^-GFP proteins. Four h collections were used, corresponding to early dividing embryos. The embryos were washed with deionized water and immediately flash frozen in liquid nitrogen and stored at −80°C before use for immunoprecipitation. Lysis and immunoprecipitation were performed similarly to S2 cell pellets except that 250 μl of embryos was used for each immunoprecipitation to saturate the beads with an excess of GFP-tagged proteins.

### On-beads protein digestion

Beads were resuspended in a digestion buffer containing 75 mM ammonium bicarbonate, pH 8.0, reduced using 10 mM DTT for 20 min and then alkylated at room temperature for 20 min in the dark with 40 mM chloroacetamide. Proteins bound to beads were digested with 1 μg of Trypsin/Lys-C mix (Promega) for 3 h at 37 °C and then supplemented with and additional amount of 150 ng Trypsin/Lys-C for overnight digestion on an end-over-end rotation apparatus at 37 °C. Peptides were isolated on C18 tips according to the manufacturer’s instructions (Thermo Fisher Scientific) and dried to completion in a SpeedVac concentrator equipped with an integrated −100°C condensation trap.

### LC-MS/MS analysis

Peptide samples were analyzed by nano LC-MS/MS on a Dionex UltiMate 3000 nanoRSLC chromatography system (Thermo Fisher Scientific / Dionex Softron GmbH, Germering, Germany) connected to an Orbitrap Fusion™ Tribrid™ mass spectrometer (Thermo Fisher Scientific, San Jose, CA, USA) equipped with a nanoelectrospray ion source. Peptides were trapped at 20 μl/min in loading solvent (2% acetonitrile, 0.05% TFA) on a 5mm × 300 μm C18 pepmap cartridge pre-column (Thermo Fisher Scientific / Dionex Softron GmbH, Germering, Germany) during 5 min. Then, the pre-column was switched online with Pepmap Acclaim column (ThermoFisher) 50 cm × 75µm internal diameter separation column and the peptides were eluted with a linear gradient from 5-40% solvent B (A: 0.1% formic acid, B: 80% acetonitrile, 0.1% formic acid) in 30 min, at 300 nl/min. Mass spectra were acquired using a data dependent acquisition mode using Thermo XCalibur software version 4.1.50. Full scan mass spectra (350 to 1800 *m/z*) were acquired in the Orbitrap using an AGC target of 4e5, a maximum injection time of 50 ms and a resolution of 120 000. Internal calibration using lock mass on the *m/z* 445.12003 siloxane ion was used. Each MS scan was followed by acquisition of fragmentation MS/MS spectra of the most intense ions for a total cycle time of 3 s (top speed mode). The selected ions were isolated using the quadrupole analyzer in a window of 1.6 *m/z* and fragmented by Higher energy Collision-induced Dissociation (HCD) with 35% of collision energy. The resulting fragments were detected by the linear ion trap in rapid scan rate with an AGC target of 1e4 and a maximum injection time of 50ms. Dynamic exclusion of previously fragmented peptides was set for a period of 20 s and a tolerance of 10 ppm.

### Database search

Mass spectra data generated by the Orbitrap™ Fusion™ Tribrid™ instrument (*.raw files) were analyzed by MaxQuant ^26^ (version 1.5.2.8) using default settings and label-free quantification (LFQ). Andromeda ^27^ was used to search the MS/MS data against a FASTA-formatted *Drosophila melanogaster* reference proteome downloaded from Uniprot ^28^ (21 923 entries) and complemented with a list of common contaminants (245 entries) maintained by MaxQuant and concatenated with the reversed version of all sequences (decoy mode). The minimum peptide length was set to 7 amino acids and trypsin was specified as the protease allowing up to two missed cleavages. Carbamidomethylation of cysteines [+57.02146 Da] was set as a fixed modification while the following mass additions were used as variable modifications: oxidation of methionine [+15.99491 Da], deamidation of asparagine and glutamine [+0.98401 Da], N-terminal and lysine acetylation [+42.01056 Da] and and pyro-glu from glutamic acid[-18.01055 Da] and glutamine [-17.02654 Da]. The reverse (decoy) and common contaminant hits were removed from the final output.

### Functional annotation

The Database for Annotation, Visualization and Integrated Discovery (DAVID) ^29, 30^ was used for protein annotation. The functional annotation tool was used to perform Gene Ontology (GO) classifications of the corresponding genes using the default settings (http://david.abcc.ncifcrf.gov/). TBCE-interacting proteins whose interaction is lost upon alanine substitution of critical amino acids in the CAP-GLY domain were selected for GO analysis. The three most significant GO categories terms for each sub-ontologies (based on GO terms *p-values*) were selected.

### Protein network modeling

Protein–protein interaction (PPI) network of TBCE was constructed using the STRING database (https://string-db.org/) ^31^. A cluster of proteins with GO assignments in the following terms were selected to generate a specific sub-network of TBCE-interacting proteins: *Biological process*: protein import into nucleus; *Cellular component*: nuclear pore; *Molecular function*: structural constituent of nuclear pore.

## Results

### dTBCE localization and fly viability depend on the presence of its conserved CAP-Gly motif

dTBCE protein has previously been isolated from taxol-stabilized MT pellets obtained from early mitotic embryos suggesting that it may have affinity for the mitotic spindle ^17, 32^. To address this question, we generated transgenic flies expressing a wild type dTBCE tagged with GFP and expressed this fusion protein under the control of a polyubiquitin promoter (Figure 1A and S1A and video 1). This transgene was functional since it successfully rescued the viability and fertility of *dtbce* mutant flies (see Methods). In addition, the expression level of dTBCE-GFP protein appeared to be similar to that of the endogenous protein (Figure S1 B, lane 2, compared with lane 1). While in early dividing embryos, dTBCE-GFP was mostly cytoplasmic, a pool of the protein was also apparent on the nuclear envelope and on the centrosome MT-aster (Figure 1A, -1:00, see also video 1). When the cells entered mitosis, dTBCE-GFP rapidly became enriched in the nuclear space right after nuclear envelope breakdown (NEBD), preceding the enrichment of tubulin in this region (Figure 1A, time 0:00, and Figure 1B, middle panel, video 1). During metaphase, dTBCE decorated the mitotic spindle region (Figure 1A, time 3:00, and Figure1B, bottom panel). During the following division, dTBCE-GFP was again detected on the MT asters and on the nuclear envelope (Figure 1A, time 16:00). dTBCE harbors a highly conserved N-terminal Cytoskeleton Associated Protein Glycine rich (CAP-Gly) motif of potential functional importance (Figure S1A). These domains are known to mediate protein/protein interactions, including binding to α-tubulin C-terminal domain ^33, 34^. It is worthy to note that a deletion of the 4 amino acids immediately downstream to this CAP-Gly motif is found in human patients suffering from hypoparathyroidy, facial dysmorphism and mental retardation (HMF, ^18^). In addition, dTBCE displays a putative C-terminal Nuclear Localization Signal (NLS). To challenge the respective importance of these two motifs, we generated several mutants of dTBCE and analyzed their live distribution as well as their ability to rescue *dtbce* mutant lethality. We first mutated the two most conserved residues of the CAP-Gly motif to disrupt its functionality (dTBCE^mutCG^, see Figure S1A). In parallel, we deleted the 4 amino acids following the CAP-Gly motif to mimic the mutation observed in HMF patients (TBCE^mutHMF^-GFP, Figure S1A). Finally, we mutated the putative NLS (TBCE^mutNLS^, Figure S1A and Methods). We noticed that both dTBCE^mutHMF^-GFP and dTBCE^mutNLS^-GFP were able to fully rescue *dtbce* mutant flies. By contrast, two independent transgenic lines expressing dTBCE^mutCG^-GFP were not able to rescue the *dtbce* lethality. Furthermore, we monitored the localization of dTBCE-GFP, dTBCE^mutCG^-GFP (Figure 1C and D), dTBCE^mutHMF^-GFP and dTBCE^mutNLS^-GFP in live embryos using a maternal driver (Figure S1C). Western blot analyses revealed that dTBCE-GFP and dTBCE^mutNLS^-GFP proteins were overexpressed at similar levels, while both dTBCE^HMF^-GFP and dTBCE^mutCG^-GFP were less abundant, suggesting that the integrity of the CAP-Gly domain may affect protein stability (Figure S2B, compare lanes 3 and 6 to lanes 4 and 5). Under these overexpression conditions, dTBCE-GFP localization pattern was similar to the one observed under moderate overexpression conditions: dTBCE-GFP was associated with the nuclear envelope and the centrosomal MT-asters during prophase, became enriched around chromosomes during prometaphase and on mitotic MTs later on (Figure1C, and video 2). Strikingly, the dTBCE^mutCG^-GFP protein showed a uniform cytoplasmic distribution, and did not associate with the nuclear envelope during interphase nor with the centrosome MT-asters or the mitotic spindle (Figure 1D, video 3). Neither the deletion adjacent to the CAP-Gly motif observed in HMF patients nor the abrogation of the putative NLS strongly impaired the wild type localization pattern of these dTBCE variants (Figure S1C and video 4 and 5). Together, this showed that while the CAP-Gly motif of dTBCE is essential for fly viability, the deletion corresponding to the mutation observed in HMF patients as well as the abrogation of the putative NLS did not significantly impair fly development or their ability to rescue the *dtbce* mutation. Altogether, these data suggest that the highly conserved amino acids of TBCE CAP-Gly motif play a key role in dTBCE function and dynamics.

**Figure 1:**
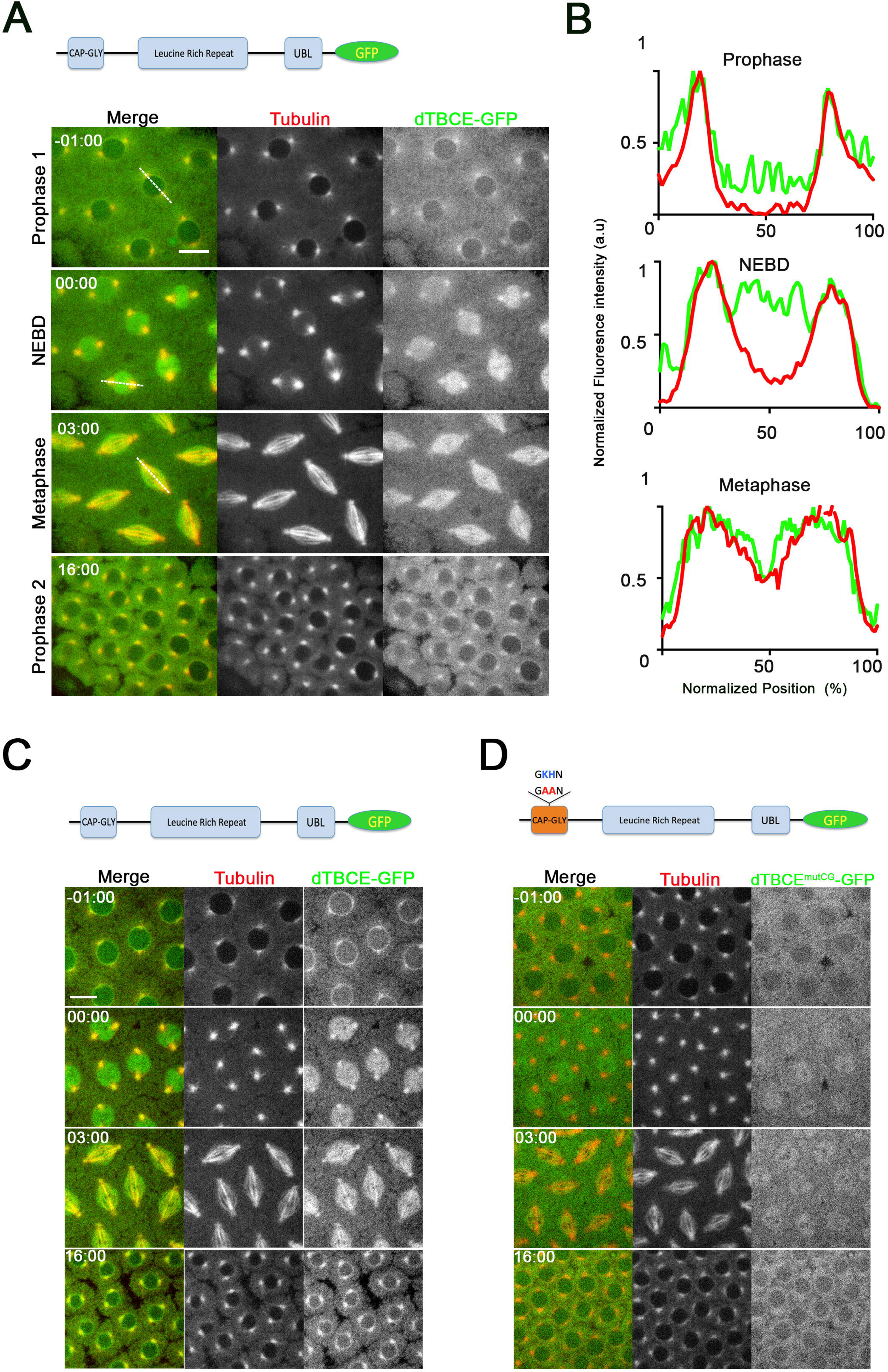
Dynamic localization of dTBCE during mitosis. A) Scheme of dTBCE domains. dTBCE harbors a N-terminal CAP-Gly motif, a central Leucine Rich Region and a C-terminal Ubiquitin-like (UBL) domain (top). Dynamics of dTBCE-GFP, expressed at endogenous levels using a poly Ubiquitin promoter (bottom, and figure S2 B). dTBCE is displayed in green in the merge panels and in monochrome in the right panels. Tubulin-RFP is displayed in red in the merge panels and in monochrome in the middle panels. Both proteins are expressed during a complete mitotic cycle of a syncytial embryo (Bottom). dTBCE-GFP localizes on the centrosome aster and on the nuclear membrane during prophase. It becomes suddenly enriched in the nuclear space during NEBD and remains in the spindle region until next cycle (see also video 1). B) Linescan intensity plots showing the normalized fluorescence intensities of dTBCE-GFP and RFP-tubulin along the dotted lines corresponding to prophase, prometaphase and metaphase stages of panel A. dTBCE-GFP is enriched in the nucleus before tubulin. C) Dynamic localization of overexpressed dTBCE-GFP using the GAL4 system and a maternal promoter (see Methods and Figure S2 B for Western blots) together with RFP-tubulin. dTBCE-GFP is displayed in green and monochrome in the right panels and tubulin-RFP is shown in red and monochrome in the middle panels. D) The conserved GKHN amino-acid sequence of the CAP-Gly motif of dTBCE was mutated into GAAN to overexpress a dTBCE^mutCG^-GFP similarly to the experiment shown in panel C. The mutant protein lost its association with MTs, the nuclear membrane, and the ability to be enriched in the nuclear space at NEBD. Scale bar: 10μm

### dTBCE associates with nuclear pore proteins and components of the Ran pathway

To deepen our molecular understanding of dTBCE function, we set out to identify dTBCE interacting partners. For this purpose, we generated S2 stable lines expressing GFP, GFP-tagged dTBCE and dTBCE^mutCG^ and performed immunoprecipitation using anti-GFP nanobodies (see methods; Figure 2A). The immunoprecipitates were subsequently subjected to proteomic analyses to identify dTBCE interacting proteins (see methods). We found that nuclear pore proteins (NUPs) were the most abundant proteins associated with dTBCE-GFP. Moreover, those associations were impaired by mutating the CAP-Gly motif (Figure 2B, C). dTBCE also displayed a strong CAP-Gly dependent association with components of the Ran pathway including α- and β-importins, as well as Ran itself, suggesting a function for dTBCE upstream or downstream of RAN (Figure 2B-C). To confirm the interaction of dTBCE with RAN *in vivo*, we prepared protein extracts from syncytial embryos expressing either GFP, dTBCE-GFP or dTBCE^mutCG^-GFP and performed immunoprecipitation experiments. These experiments confirmed that dTBCE-GFP interacts with Ran as well as with α-tubulin and demonstrated that the strength of these interactions were CAP-Gly dependent in early embryos, similarly to S2 cells (Figure 2D). The gene ontology classification of dTBCE interacting protein whose interaction was lost upon mutation of the CAP-Gly domain highlights the fact that this protein is located at the boundary between the nuclear space and the cytoplasm and can interact with nucleocytoplasmic transport proteins and with microtubule (Figure 2E).

**Figure 2:**
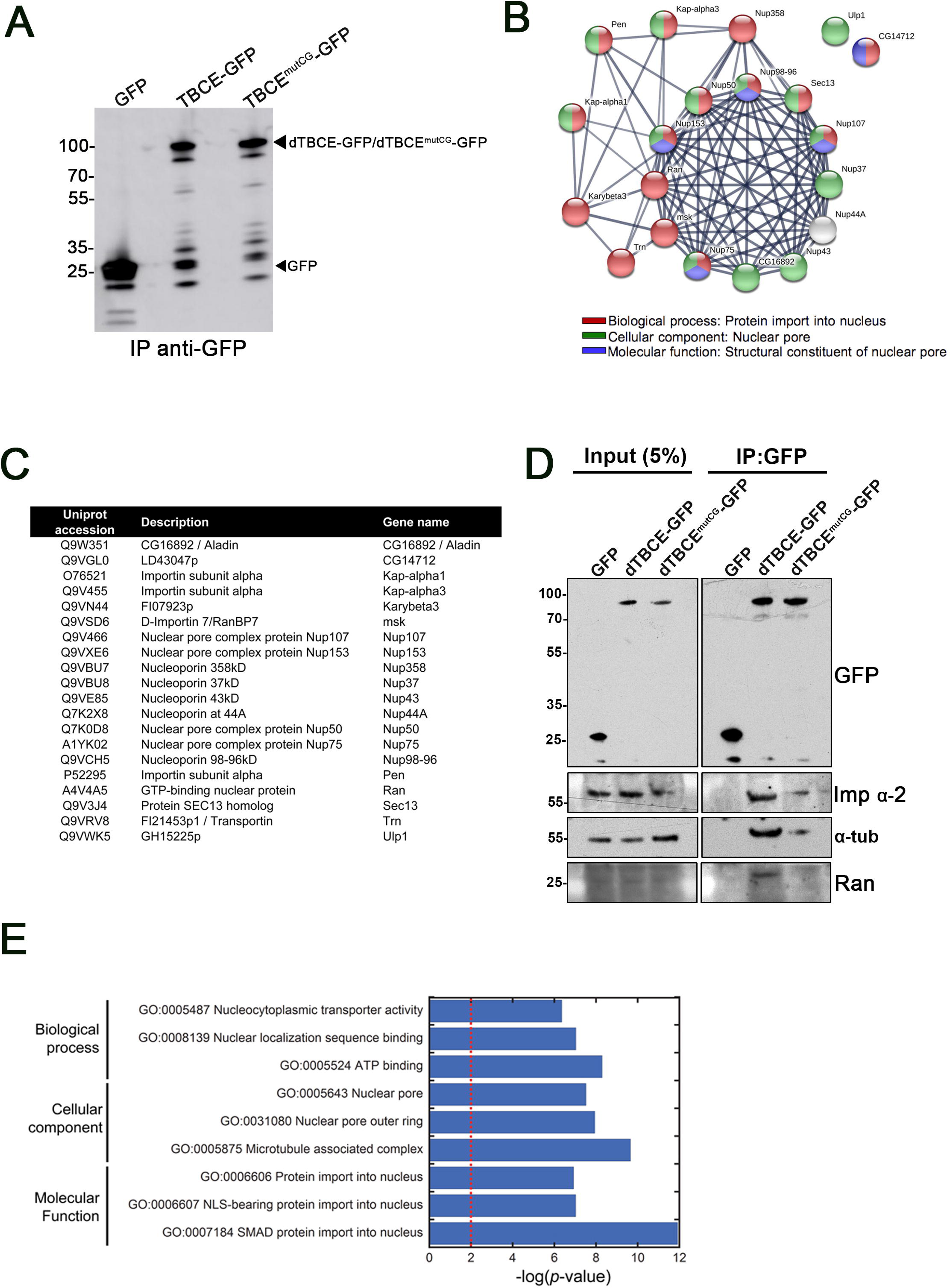
Interaction of dTBCE with Nuclar Pore Proteins and components of the RAN pathway. A) Lysates of stable S2 cell lines expressing GFP, dTBCE-GFP, and the non-functional CAP-Gly mutant dTBCE^mutCG^-GFP, were collected and subjected to immunoprecipitation with anti-GFP nanobodies (see Methods). A fraction of the immunoprecipitation reaction (5%) was analyzed by Western blot using an anti-GFP antibody, and the remaining fraction was analyzed by mass spectrometry to identify dTBCE-interacting proteins. B) Interaction network of dTBCE interacting proteins. The cluster was generated by STRING (see Methods). C) List of interacting proteins used for the scheme shown in B. D) Anti-GFP Immunoprecipitation experiments were performed from embryo extracts expressing either GFP, dTBCE-GFP or dTBCE^mutCG^-GFP. The samples were subsequently analyzed by Western blot to confirm the presence of GFP, GFP-tagged dTBCE variants, α-tubulin and Ran. E) Gene Ontology classification of dTBCE-interacting proteins whose interaction is lost upon substitution of critical amino acids in the CAP-Gly domain. The graph summarizes the three most significant GO categories terms for each sub-ontologies. The significance of enrichment is expressed as –log(p-value). The red dashed line indicates a 0.01 p-value threshold.

### Lowering dTBCE levels impairs brain growth and triggers spindle assembly defects associated with a SAC-dependent mitotic delay

dTBCE was initially described as a gene potentially involved in mitotic spindle assembly of larval brain Nbs ^17^. The localization pattern of dTBCE and its interaction with Ran, a master player in the nucleation of spindle MTs around chromatin, prompted us to carefully examine dTBCE-dependent mitotic spindle assembly defects. We therefore knocked down in the fly central nervous system dTBCE using a two-days treatment with one or two independent *dTBCE* specific dsRNAs (see Methods). While the knock down of the tubulin-binding co-factor dTBCD leads to a decrease in tubulin levels ^35^, we found that RNAi-driven down regulation of dTBCE did not affect either α- or β-tubulin protein levels (Figure 3A). Strikingly, dTBCE-depleted brain lobes were smaller than controls, indicating neural tissue growth defects (Figure 3B, *n*>50 brains). In brain squashes analyses, the chromosomes of dTBCE-depleted brains cells were frequently over condensed likely indicating a mitotic delay (Figure 3C). In agreement with these observations, mitotic indices analyses revealed that knock down of dTBCE with a single or a combination of two dsRNAs triggered respectively a significant 2 and 4 times elevation of the mitotic indices (Figure 3D). Subsequent fixation of brain tissues and staining for tubulin revealed that dTBCE-depleted Nbs metaphases exhibited a strong decrease in MT density within the spindle region (Figure 3E, compare left to right). Moreover, these Nbs exhibited significantly shorter spindles compared to control Nbs (Figure 3F). Analyses of dTBCE-depleted Nbs expressing β-tubulin-GFP by live microscopy confirmed the decrease in tubulin signal within the middle region of the metaphase spindles (Figure 3G, bottom panels, time 3:00 and 5:00, red arrowheads). Moreover, while mitotic duration was 5 min in control Nbs, knock down of dTBCE by a single or a double RNAi treatment led to a significant increase of the mitotic timing (Figure 3H). Mitotic delays and spindle abnormalities are frequently associated with Spindle Assembly Checkpoint (SAC) activation. To challenge the contribution of the SAC to the mitotic delay observed in dTBCE-deficient cells, we first analyzed the mitotic indices of mitotic Nbs subjected to RNAi for the spindle assembly checkpoint protein Mad2, dTBCE or both (Figure S2A-B). We noticed that double Mad2 and dTBCE knock down led to the disappearance of mitotic cells with over condensed chromosomes and the appearance of aneuploid cells (Figure S2A, red arrowhead). In addition, the mitotic index, which was 2.39 ± 0.35 % in dTBCE knock down neural tissues, dropped to 0.75 ± 0.11 % when both dTBCE and Mad2 were knocked down (*P*< 0.0001, Figure S2B). By analyzing the mitotic timing of *dTBCE* RNAi treated Nbs in the presence or absence of Mad2 dsRNA (Figure S2C-D), we found that knock down of both Mad2 and dTBCE led to a complete rescue of mitotic timing. Finally, in fixed preparations, we frequently detected lagging chromosomes in anaphase cells when both Mad2 and dTBCE were knock down, indicating that the SAC is required to maintain genome stability in cells with low levels of dTBCE, in agreement with the presence of aneuploid cells in that context (Figure S2 A and E). Altogether, our data indicates that dTBCE is essential for proper mitotic spindle assembly in neural stem cells, and is required for normal larval brain growth.

**Figure 3:**
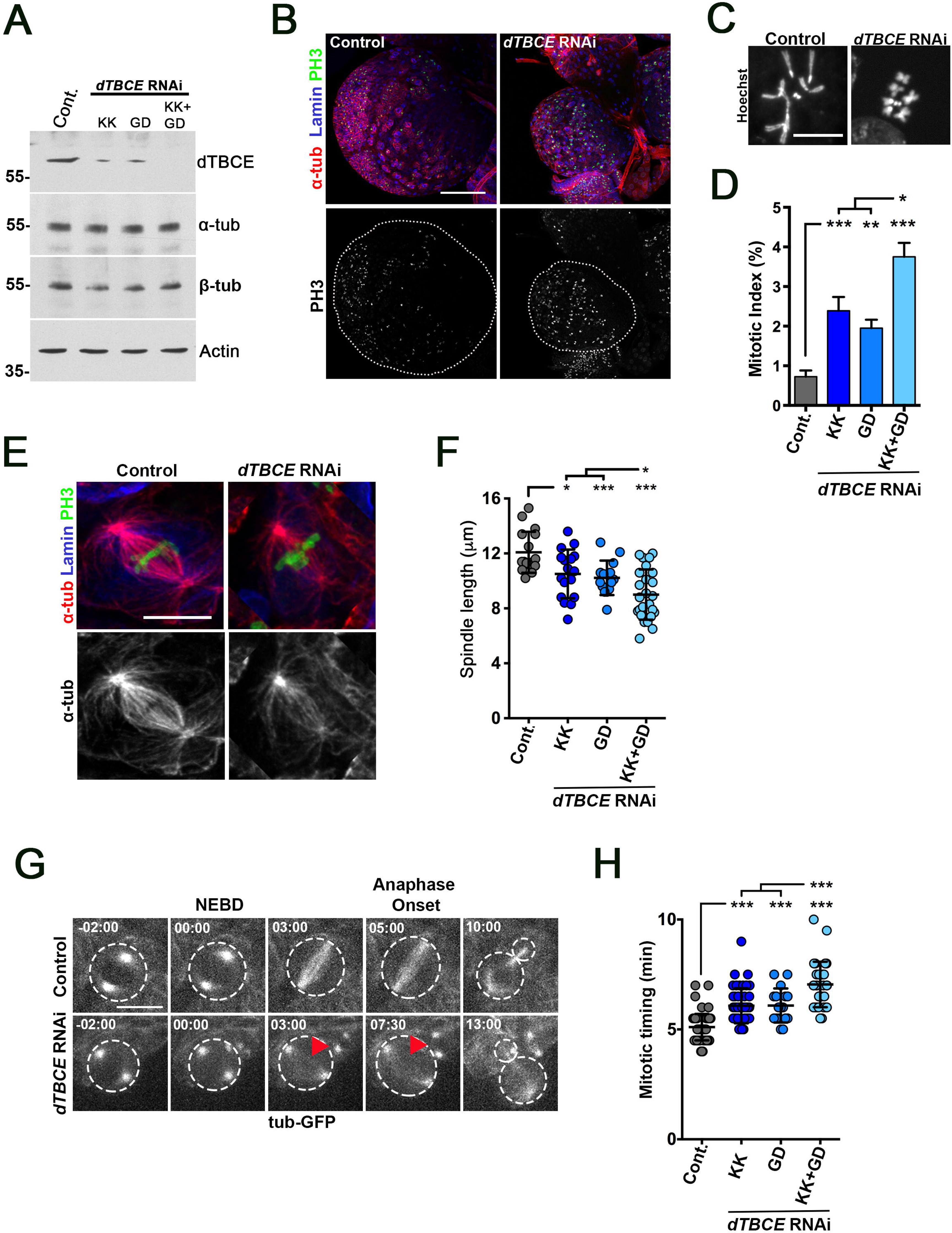
Importance of dTBCE for neural tissue development and mitotic spindle assembly in *Drosophila* neuroblast. A) Analysis of dTBCE, α-tubulin, β-tubulin and actin protein levels in fly neural tissues in control or *dTBCE* RNAi samples. Two RNAi lines were used (named KK and GD, see Methods), alone or in combination to drive more efficient knockdown of dTBCE. B) Overview of optic lobes of control (left panels) and *dTBCE* double RNAi larval brains (right panels) stained for tubulin (red), lamin (blue) and phospho-Histone H3 (PH3, green and lower panels in monochrome). The lobe contour is visualized with a dotted line in the lower panels. *dTBCE* RNAi lobes display growth defects compared to controls. Scale bar: 100 μm. C) Mitoses of squashed preparations of mitotic cells from control (left) or *dTBCE* dsRNAs treated brains (right). The chromosomes are often hyper condensed in *dTBCE* RNAi brains. D) Histogram showing the mitotic index (± s.d.) in control brain tissues: 0.73 ± 0.16 (*n*=13095, 3 brains) or in *dTBCE* RNAi brains: KK 2.39 ± 0.35 (*n*=11535, 4 brains), GD: 1.95 ± 0.21 (*n*=6329, 2 brains) and KK+GD: 3.75 ± 0.35 (n=6319, *2* brains). E) Metaphase in control and *dTBCE* RNAi Nbs. Microtubules are in red (and monochrome in the lower panels), DNA in green and lamin in blue. Note the apparent decrease in mitotic spindle MT density in *dTBCE* RNAi (right) compared to the control (left). F) Quantification of spindle length in control (12.1 ± 1.5μm *n*=17) or *dTBCE* RNAi (KK: 10.5 ± 1.8μm *n*=17; GD: 10.2 ± 1.3μm *n*=14; KK+GD: 9.0±1.8μm, *n*=25). G) Mitosis in WT (top panels) or in *dTBCE*-depleted live Nbs (bottom panels) expressing β-tubulin-GFP. Note the apparent lower MT density (red triangles) and the delay between NEBD and anaphase onset following dTBCE depletion with one or two dsRNAs. Time is min:s. Scale bars are 10μm for panels C, E and G. H) Dot Plot (± s.d.) showing mitosis duration in control (5.1 ± 0.6 min, *n*=67) and *dTBCE* RNAi Nbs (KK: 6.1 ± 0.8 min, *n*=45; GD: 6.1 ± 0.8 min, *n*=17; KK+GD: 7.0 ± 1.0 min, *n*=32). *: *P*<0.05, **: *P*<0.01, ***: *P*<0.001, Wilcoxon test.

### dTBCE is essential for the polymerization of mitotic spindle MTs

To investigate the origin of the low MT density observed in the spindle region of dTBCE deficient cells, we first analyzed MT regrowth at 25°C following cold-induced full MT depolymerization (Figure 4A) as described before ^17^. In control cells, a strong MT regrowth was observed around the chromosomes (red arrowhead) and at the centrosomes after 30 s. In addition, the mitotic spindles displayed a normal bipolar structure after only 90 s (Figure 4B, left). By contrast, in dTBCE knock down cells, a severe impairment of MT nucleation was observed on the kinetochore and the spindle region after 30 s at 25°C while nucleation at the centrosome appeared to remain normal (Figure 4B right, middle panel). After 90 s, *dTBCE* RNAi mitotic Nbs retained this apparent low MT density on the spindle region (Figure 4B, right). We next decided to analyze and quantify MT nucleation in space and time in intact metaphase flattened Nbs by live tracking the MT plus end binding protein EB1-GFP (Figure 4C-D). In controls, time projections and kymographs revealed that MT nucleation and elongation occurred at the centrosomes (black arrows), and also very strongly in the spindle region itself (Figure 4C, left and video 5). In dTBCE knock down Nbs, MT nucleation was similar to control Nbs at the centrosome but was clearly diminished at the spindle region, suggesting that this process was impaired around chromatin (Figure 4C, right and video 6). Quantitation of the EB1-GFP intensity signal ratio between the chromosome region and the centrosome revealed a 50 % decrease between control and dTBCE-depleted cells (See Methods; Figure 4D, right; *P*<0.0001), while EB1-GFP protein levels remained stable in both control and *dTBCE* RNAi treated cells (Figure 4E). Together, these results reveal that MT nucleation around chromatin relies on dTBCE.

**Figure 4:**
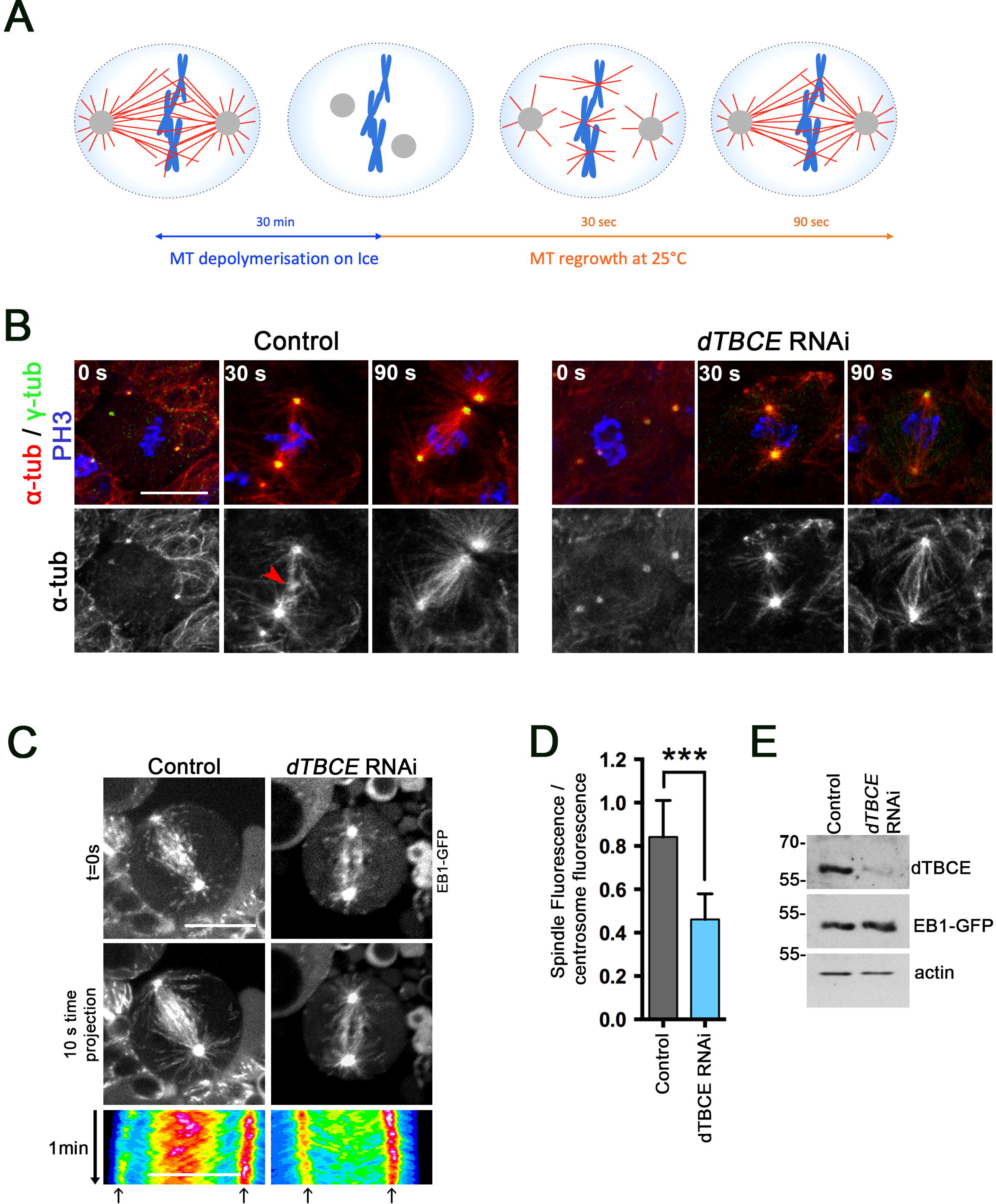
Contribution of dTBCE to microtubules nucleation around chromatin. A) Scheme illustrating the MT depolymerization and regrowth assay. Fly brains were dissected in Schneider media and incubated on ice for 30 min to induce total MT depolymerization. MT regrowth was induced by a temperature shift to 25°C. Brains were fixed after 30 s, a step at which MT nucleation can be visualized at the centrosomes and around chromatin. 90 s following return to 25°C, a bipolar spindle is formed. Centrosomes are displayed in grey, MTs in red and DNA in blue. B) Representative images of control (top panels) and *dTBCE* RNAi (bottom panels) mitotic Nbs during microtubule regrowth after cold-induced MT depolymerization (see Methods). Samples were analyzed immediately after depolymerization (0 s, left panel), 30 s (middle panel) and 90 s (right panel) after return to 25° C. The red arrowhead shows MT nucleation around chromatin at 30 s, which is absent when *dTBCE* is depleted. MTs are shown in red (top panels and monochrome in the lower panels), DNA in blue and γ-tubulin in green. Scale bar: 10μm. C) Control (left) and *dTBCE* RNAi (right) metaphase Nbs expressing EB1-GFP were imaged by time-lapse spinning disk confocal microscopy. A single frame is presented at the top panel and a 20-time points projection (10 s) is displayed in the middle panels. The bottom panel shows kymographs of microtubule plus ends intensity along the duration of the time-lapse acquisition (1 min) between the two spindle poles (LUT: dark blue to red). The arrows point the position of the centrosomes. Comet velocity in controls (16.75μm/min, *n*=10 cells) and in *dTBCE* RNAi treated cells (16.11μm/min, *n*=12) was similar (*P*=0.46, Wilcoxon test) measured using the U-track software. A severe decrease in the nucleation of MTs is observed around the metaphase plate. D) Histogram showing the mean EB1-GFP fluorescence intensity ratio (a.u.) between chromatin and the centrosome in control (0.84 ± 0.16, *n*=7) and *dTBCE* RNAi metaphase neuroblasts (0.46 ± 0.14, *n*=14). ***: *P*<0.001. E) Western blot analyses showing dTBCE (top), EB1-GFP (using an anti-GFP antibody, middle), and actin protein levels (bottom) in control or in dTBCE depleted-brains tissues.

### Ran and dTBCE are required for tubulin enrichment in the nuclear space during mitosis

In agreement with previous studies ^36–38^, our data support the direct interaction of dTBCE with tubulin (Figure 2D). In addition, we found that dTBCE interacts with the Ran GTPase and both proteins are required for the nucleation of MTs around chromatin. Strikingly, the sudden enrichment of dTBCE-GFP in the nuclear space, shortly before tubulin recruitment, prompted us to determine the dependency of these two events. Such a tubulin enrichment process is known to take place in the fly and worm embryos ^13, 15^. Interestingly, this event was shown to be under the control of Ran in *C. elegans* ^13^. We therefore asked whether a tubulin enrichment process in the nuclear space was present in the fly Nbs. Since tubulin signal from MTs nucleated at the centrosomes would likely mask tubulin import in the nuclear space, we decided to depolymerize the MT network by incubating the tissue with colchicine (see methods, Figure 5A, and compare Figure S3A and B). Under these conditions, we found that the maximum tubulin intensity was reached after 60 s following NEBD (Figure 5B top panels and Figure 5C, grey curve). In Ran RNAi Nbs, the intensity of the tubulin signal slowly and progressively reached equilibrium between the nuclear and the cytoplasmic pool (Figure 5B, lower panels and Figure 5D, orange curve). This indicates that tubulin has a poor ability to reach the nuclear space in the absence of Ran. When dTBCE was knocked down, tubulin never became enriched in the nuclear space (Figure 5B, middle panels, and Figure 5C, blue curve). To better understand the limited ability of tubulin to reach the nuclear space, we analyzed mitotic spindle compartmentalization during mitosis by live microscopy using several fluorescent membrane markers (Figure 6). The nuclear pore protein Nup107-RFP rapidly disappeared at mitotic entry and reappeared during late telophase (Figure 6A). Analysis of lamin-GFP dynamics indicated that the nuclear membrane persisted around the mitotic apparatus, since 50 % of the total lamin-GFP signal was still present during metaphase (Figure 6B). Finally, monitoring total membranes, using a CD8-GFP reporter showed that the Nb mitotic spindle was embedded in membranes all along cell division (Figure 6C). Therefore, as previously suggested for human and *Drosophila* S2 cells ^14, 39^, nuclear and cytoplasmic membranes reside around the mitotic apparatus, which account for the limited ability of tubulin to easily enter the nuclear space. Altogether, these experiments reveal that the ability of tubulin to reach the nuclear space is restricted and that active enrichment of tubulin depends on both dTBCE and Ran (Figure 5E).

**Figure 5:**
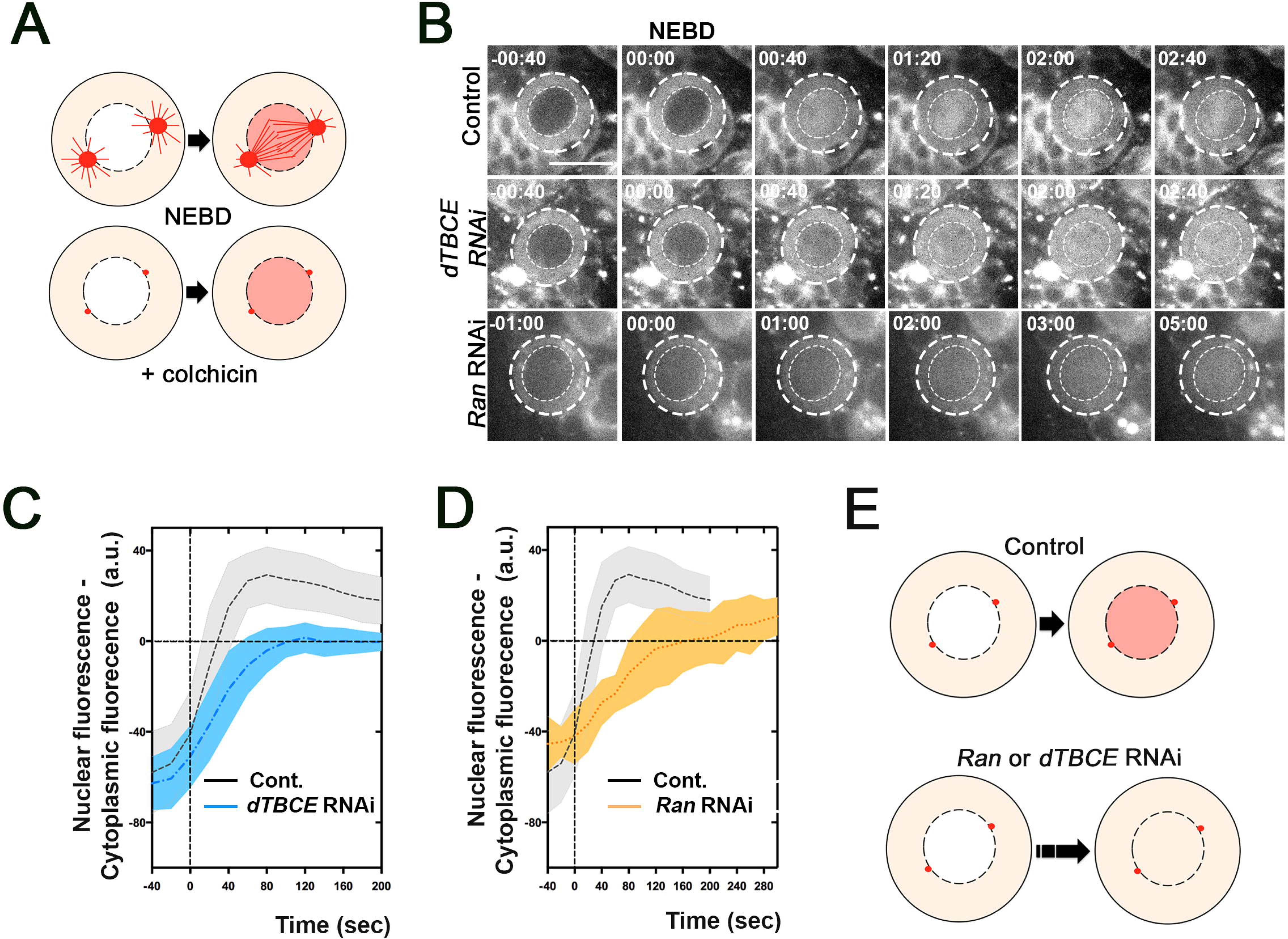
Role of dTBCE in enriching tubulin in the nuclear space. A) Scheme showing tubulin import in the nuclear space after NEBD in the presence (top) or absence of MTs (bottom, + colchicine). Depolymerizing conditions were used to get rid of tubulin signal from MTs (spindle and centrosomes), and thus allow an accurate quantification of the tubulin signal in the nuclear compartment. B) Selected live imaging time points of control (top), dTBCE (middle) and Ran-knock down Nbs (bottom) expressing mCherry-α-tubulin during early mitosis after incubation with 20 μM colchicine to depolymerize the MT network. The time is indicated as min:s. Scale bar is 10 μm. Note the accumulation of tubulin in the nuclear region just after NEBD in the control at time 00:40. C) Graphs (± s.d.) displaying the difference in signal intensity between the nucleus and the cytoplasm among time (0:00 corresponds to NEBD) in control (grey) or dTBCE knock down cells. D) Curves (± s.d.) displaying the difference in signal intensity between the nucleus and the cytoplasm among time (0:00 corresponds to NEBD) in control (grey) or Ran knock down cells (orange). Note the low diffusion of tubulin in the nuclear space when Ran or dTBCE are depleted. The control is the same in C and D. E) Scheme recapitulating tubulin enrichment in the nuclear space at NEBD in control (top) and Nbs depleted for dTBCE or Ran Nbs (bottom).

**Figure 6:**
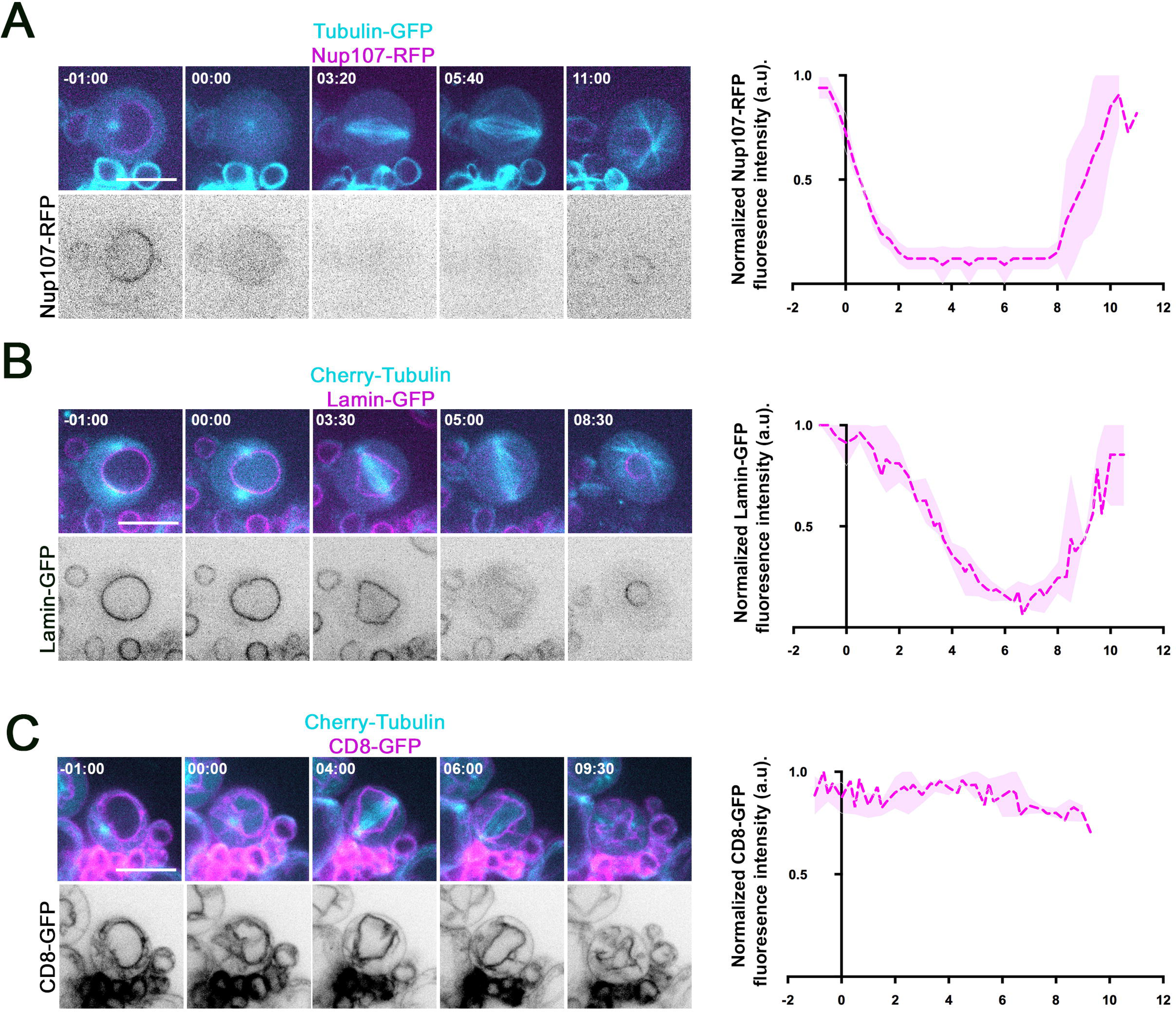
Neuroblast mitotic spindle compartmentalization during mitosis. A) Left: Selected frames of a flattened Nb expressing β-tubulin-GFP (cyan) and Nup107-RFP, a nuclear pore protein (magenta in the merged image, grey in the bottom line). Right: Quantification of the normalized average fluorescence intensity (± s.d.) of Nup107-RFP on the nuclear membrane during mitosis. Nup107-RFP rapidly disappears at NEBD. B) Left: Selected frames of a flattened Nb expressing mCherry-α-tubulin (cyan) and lamin-GFP, a nuclear membrane protein (magenta in the merged images, grey in the bottom). The lamin-GFP labeled membrane surrounds the mitotic spindle from NEBD till anaphase onset. Right: Quantification of the normalized average fluorescence intensity (± s.d.) of lamin-GFP around the nucleus/mitotic apparatus during mitosis. The lamin-GFP signal slowly disappears after NEBD until anaphase onset. C) Left: Selected frames of a flattened Nb expressing mCherry-α-tubulin (cyan) and CD8-GFP to visualize all the cell membranes during cell division (magenta in the merged images, grey in the bottom). Left: quantification of the normalized average fluorescence intensity (± s.d.) of CD8-GFP around the nucleus/mitotic apparatus during mitosis. CD8-GFP remains at a high level around the mitotic apparatus during cell division. T0 corresponds to NEBD, *n*=3 cells for each genotype. Scale bars: 10μm.

### dTBCE dynamically traps tubulin in the nuclear space after NEBD

To investigate how dTBCE could be involved in tubulin enrichment in the nuclear space, we first monitored the dynamics of both Cherry-α-tubulin and dTBCE-GFP in this region. Our live localization studies in live embryos showed that dTBCE was first enriched in the nuclear space at NEBD (Figure 1A and video 1). To confirm these observations, we treated Nbs with colchicine and found that dTBCE was enriched before tubulin in the nuclear space, supporting the hypothesis that dTBCE is not traveling along with tubulin, but is rather imported in the nucleus by Ran with other proteins such as Nups to form there a tubulin trap (Figure S3B). To verify the ability of dTBCE to trap tubulin, we immunoprecipitated and immobilized either GFP, dTBCE-GFP, or dTBCE^mutCG^-GFP on beads (see Methods) and assayed the ability of these beads to trap purified tubulin-rhodamin (Figure 7A-C). We found that GFP, dTBCE-GFP, or dTBCE^mutCG^-GFP were present at similar levels at the surface of the beads (Figure 7A and B). However, only dTBCE-GFP coated beads were able to trap pure tubulin (Figure 7A and C). FRAP analyses revealed a continuous exchange between bound and free tubulin in the photo bleached region (Figure 7D and E), indicating that the tubulin pool at the surface of the beads was reversibly bound. These experiments support a mechanism through which dTBCE is imported in the nuclear space to create a dynamic trap that leads to tubulin enrichment prior to spindle assembly (Figure 7F).

**Figure 7:**
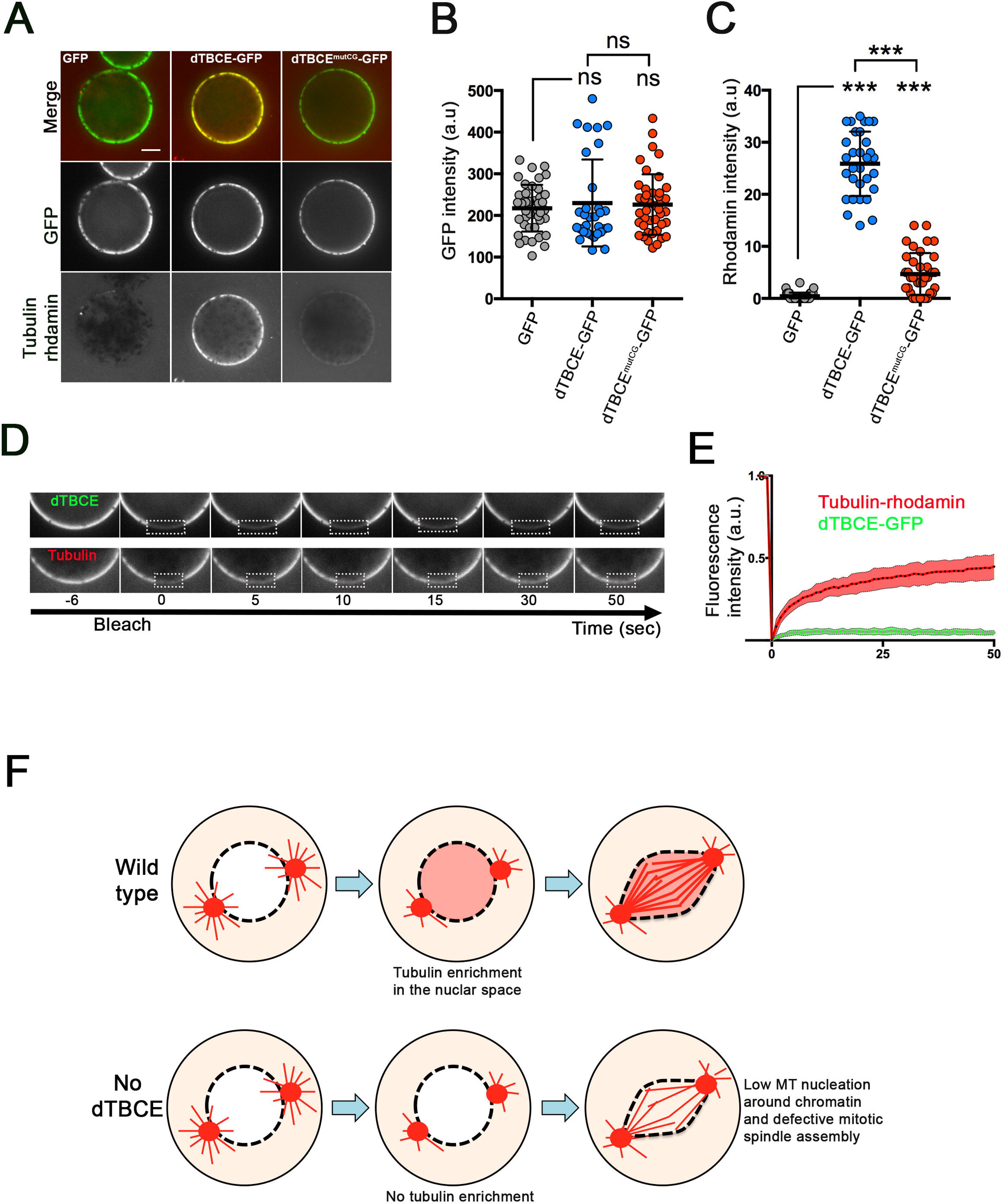
Tubulin trapping by dTBCE. A) GFP, dTBCE-GFP and dTBCE^mutCG^-GFP were immunoprecipitated from S2 cells and bound to GFP-trap beads (green, top panels). Beads were subsequently incubated with rhodamine-conjugated tubulin and GTP. The beads were then imaged by spinning disk-confocal microscopy (see Methods). B) Dot plot (± s.d.) showing the mean fluorescence intensity (a.u.) of each sample in the GFP channel (left, GFP: 217.4 ± 56.0, *n*=42; TBCE-GFP: 229.9 ± 104.4, *n*=31 and TBCE^mutCG^-GFP 226.2 ± 76.6, *n*=42). C) Dot plot (± s.d.) showing the mean fluorescence intensity (a.u.) of each sample in the tubulin-rhodamine channel (right, GFP: 0.4 ± 0.7, *n*=42; dTBCE-GFP: 25.9 ± 6.2, *n*=31 and dTBCE^mutCG^-GFP 4.7 ± 4.0, *n*=42). ***: *P*<0.001 (Wilcoxon test). Tubulin-rhodamine is present at the surface of dTBCE-GFP beads, absent from GFP beads and strongly reduced when the CAP-Gly motif of dTBCE was mutated. D) FRAP experiment, showing the recovery after photo bleaching (FRAP) of a small region (indicated by a dotted rectangle) at the surface of dTBCE-GFP beads incubated with tubulin-rhodamine, in the green and red channel respectively. A recovery of tubulin fluorescence is observed indicating an exchange between the dTBCE-interacting tubulin and the free tubulin pool. E) Quantification of the GFP (green curve, *n*=4) and the tubulin-rhodamine (red curve, *n*=9) intensities (± s.d) in the photo bleached regions beads for the GFP and). F) Possible model for dTBCE-dependent tubulin import in the compartmentalized nuclear space during early mitosis. MTs are displayed as red lines and the levels of tubulins are reflected by the orange intensities. In wild type *Drosophila* cells (top), dTBCE and nuclear pore components are incorporated in the nuclear space to create a tubulin trap. Tubulin (red) is then enriched through the dTBCE-CAP-Gly motif in the future spindle region. This enrichment of tubulin promotes nucleation of MTs around chromosomes. When dTBCE is knock down (bottom), tubulin fails to be efficiently enriched in the nuclear space. The low tubulin concentration prevents sufficient MTs enough nucleation around chromatin impairing mitotic spindle assembly. Cytoplasmic and nuclear membranes that surround the mitotic apparatus (dotted line) persist at least until metaphase.

## Discussion

Although many differences exists between model systems, several decades of *in vitro* and *in vivo* genetic studies have shown in eukaryotic systems that tubulin binding co-chaperones contribute to the dynamics of MTs ^16, 40^. In agreement with this hypothesis, it has been shown that manipulation of chaperone levels in *Drosophila* and mammalian cells triggers defective MT polymerization caused by an overall depletion of tubulin ^18, 19, 35, 41–43^. Interestingly, our experiments reveal that low levels of dTBCE are sufficient to sustain normal tubulin dimer formation, in agreement with previous genetic studies in Drosophila and budding yeast ^19, 44^. Moreover, our proteomic analyses with a functional dTBCE-GFP transgenic protein did not reveal any detectable interaction with other tubulin-binding co-chaperones, in agreement with a previous report in *Drosophila* ^41^. Therefore, although tubulin co-chaperones can be found in a common protein complex with tubulins when produced and manipulated *in vitro* ^16, 37^, it is possible that the chaperone complex is unstable *in vivo*, and assembled only under particular stress conditions as suggested by deletion studies in budding yeast ^44^. Alternatively, we cannot rule out that this complex is present in too low amounts to be detected in our proteomic analyses. Along this line, dTBCE may have diversified its functions, during evolution of genomes in addition to its initial role as a tubulin chaperone. Congruently, analyses of dTBCE interacting proteins revealed robust physical interactions with Nuclear Pore Proteins and components of the Ran pathway (Figure 2). Another new and unexpected feature of dTBCE came from the study of its dynamic localization in Nbs and early embryos of *Drosophila*. Mammalian TBCE is a cytoplasmic protein (unpublished result and ^43^). We here show that dTBCE is more specifically present on the centrosomal MTs and associated with the nuclear membrane during interphase, before being suddenly enriched in the nucleus at the time of NEBD (Video 1). Interestingly, the conserved CAP-Gly motif of dTBCE is necessary to restore the viability of null *dtbce* mutants, to promote the correct dynamics of dTBCE, to mediate the interaction with NUPs, the Ran machinery, and to interact with tubulin. Interestingly enough, dTBCE knockdown leads to a loss of MT nucleation specifically around chromatin, leading to the formation of shorter mitotic spindles harboring low MT density. This mitotic phenotype and the physical interaction of dTBCE with Ran, a key regulator of chromatin-mediated MT nucleation, is interesting because in *C. elegans* embryo, tubulin enrichment in the nuclear space is regulated by Ran, and is proposed to participate in mitotic spindle assembly ^13^. This process has probably been underestimated since it is difficult to visualize “soluble” tubulin enrichment at NEBD, when large microtubule organizing centers nucleating numerous MTs and concentrating high levels of tubulin are present. However, under MT depolymerizing conditions, tubulin concentration in the nuclear space is visible during NEBD in *D. melanogaster* and *C. elegans* embryos ^13, 15^. We have been able to confirm that a similar tubulin enrichment pathway is also present in the stem cells of the fly central nervous system. Lowering dTBCE level impairs this process, resulting in a subsequent weaker MT nucleation around chromatin and the generation of spindle assembly defects. How Ran and dTBCE proteins cooperate to achieve tubulin import remains to be explored in details. However, live observation of tubulin and dTBCE dynamics in NBs and embryos have shown that dTBCE is enriched first in the nuclear space, suggesting that the two proteins do not “travel together”. Therefore, we propose that following NEBD, dTBCE is imported to generate a dTBCE-rich environment that favors, on its own or together with NUPs, subsequent dynamic tubulin trapping through its CAP-Gly motif. Our results show that dTBCE-GFP beads (also containing NUPs and Ran) are able to concentrate and exchange tubulin. Such enrichment and tubulin turnover are likely essential to provide a reservoir of free tubulin available for polymerization of spindle MTs. Of note, we find that dTBCE-GFP beads are not sufficient to trigger MT nucleation, which suggest that other factors are needed to reconstitute a full nucleation pathway. A key result of our study is the physical interaction of dTBCE with members of the RAN pathway which are important players of mitosis and meiosis in many systems ^6^. Similarly to dTBCE, Ran also controls tubulin import in the nucleus (this work, ^13^). Because dTBCE does not harbor a functional nuclear localization signal, we therefore believe that dTBCE is not directly imported in the nucleus by the Ran machinery and cannot be considered as a “true” Spindle Assembly Factor that displays a NLS. We hypothesize that Ran would be a leader in reorganizing the nuclear space to create an environment conducive to tubulin enrichment for the nucleation and polymerization of spindle MTs (this paper ^13, 45, 46^). We envision three advantages in accumulating tubulin in the spindle region. First of all, fly cells are described in textbooks to display a semi-open mitosis. This vague notion leaves the reader with the idea that molecules can nevertheless freely travel between the spindle and the cytoplasm through a fenestrated membrane. However, when monitoring nuclear membrane dynamics in live cells, we found that lamin was retained around the mitotic spindle until metaphase. Moreover, in agreement with studies in S2 cells ^39^, we also found that cell membranes surround the mitotic apparatus until late telophase. Therefore, this notion of semi-open mitosis should be revisited, since the compartmentalization of the mitotic apparatus during cell division likely restrains the diffusion of tubulin. Second, we think that accumulating tubulin in the nascent spindle region is particularly advantageous for cells with nuclei of large volume, such as Nbs. A simple prediction of such a high nucleus to cell size ratio is the dilution of tubulin concentration following NEBD, leading to defective MT nucleation. Finally, we find that the nuclear space is not an environment that favors fast diffusion of tubulin when either Ran or dTBCE are absent (Figure 5C and D) in agreement with similar observations made in the one cell *C.elegans* embryo ^13^. Consequently, in absence of an active tubulin enrichment mechanism at NEBD, fly cells are exposed to constraints that are limiting MT nucleation around chromatin. In agreement with tubulin import being a key event, preventing this process delayed kinetochore capture and triggered activation of the SAC. Suppression of the SAC in dTBCE knocked-down cells leads to chromosome segregation defects, suggesting that tubulin import is crucial for genome stability. Interestingly, we noticed that dTBCE might also be used for MT nucleation in other subcellular compartments. Indeed, MmTBCE is recruited to the Golgi apparatus and is required for nucleation of Golgi-derived MTs. However, it is not clear whether MmTBCE triggers local tubulin concentration to mediate MT nucleation ^47, 48^. Moreover, it has been shown that tubulin enrichment at the centrosome by MAPs may be an essential mechanism to promote nucleation of MTs ^11, 12^. It will thus be interesting to investigate whether local increase in tubulin concentration by tubulin-interacting proteins is a general and conserved mechanism involved in the spatiotemporal regulation of MTs.

## Acknowledgments

This work was supported by the Ligue Nationale contre le Cancer, the Fondation ARC pour la Recherche Contre le Cancer, the University of Rennes 1 and the Centre National de la Recherche Scientifique. MM is fellow of the Region Bretagne, the Ligue Nationale contre le Cancer and the Fondation ARC pour la Recherche Contre le Cancer. EG is a post-doctoral fellow of the Fondation pour la Recherche Médicale. We are grateful to Roger Karess and Valérie Doye (Institut Jacques Monod Paris), Yong Zhang and Shan Jin (Institute of Genetics and developmental Biology, Beijing) for the kind gifts of fly stocks, DNA constructs, and antibodies. We would like to thank Sébastien Huet, Stéphanie Dutertre and Xavier Pinson for their help using the microscopes of the Microscopy Rennes imaging center platform.

---

**Figure S1: Western blot analysis and localization of dTBCE variants *in vivo*** A) Scheme of the different dTBCE variants used for live imaging and rescue experiments. Wild type dTBCE (top) harbors a N-terminal CAP-Gly motif, a central Leucine Rich Region and an ubiquitin-like domain (UBL) containing a predicted weak Nuclear Localization Sequence (NLS). The dTBCE^mutCG^ (mutCG: mutated CAP-Gly) variant was generated by replacement of the most conserved GKHN sequence into GAAN (second from the top). A deletion of 4 amino acids downstream the GKHN motif was also created to mimic the mutation found in HMF human patients and named dTBCE^HMF^ variant (third from the top). Finally, the putative C-terminal NLS motif (MKRPRL) was mutated into (MAAPAL) to generate a NLS defective protein dTBCE^mutNLS^ (bottom). All these proteins were expressed with a C-terminal GFP tag expressed under the control of a poly ubiquitin promoter (dTBCE-GFP only) or using the UAS-GAL4 system (for dTBCE-GFP, dTBCE^mutNLS^-GFP, dTBCE^mutCG^-GFP and dTBCE^HMF^-GFP). B) Western blot analysis of dTBCE expression in embryos using an anti-dTBCE antibody (top) or an anti-actin antibody (bottom). Lane1: control embryo extract, Lane 2: dTBCE-GFP embryo extracts in which expression of dTBCE-GFP is under the control of a poly ubiquitin promoter, Lane 3 to 6: dTBCE-GFP, dTBCE^mutCG^-GFP, dTBCE^HMF^-GFP, and dTBCE^mutNLS^-GFP extracts in which expression of the corresponding proteins was driven by the UAS/GAL4 system using a maternal driver (v32c-Gal4). C) Localization of dTBCE^HMF^-GFP (top) and dTBCE^mutNLS^-GFP (bottom) in early dividing embryos. Time is min:s. Scale bar: 10 μm.

**Figure S2. dTBCE and Mad2 co-depletion shortens mitotic duration and triggers severe chromosome segregation defects during anaphase.** A) Mitotic figures of squashed preparations from neural tissues. Control (top left), dTBCE-depleted (top right), Mad2-depleted (bottom left) and dTBCE+Mad2-codepleted mitotic cells (bottom right) are shown. The arrowhead points an additional X chromosome. B) Histogram showing the mitotic indices (± s.d.) in control brain tissues: 0.73 ± 0.16 (*n*=13095, 3 brains), in *dTBCE* RNAi brains: KK 2.39 ± 0.35 (*n*=11535, 4 brains), in *Mad2* RNAi brains: 0.73 ± 0.07 (*n*=7655, 3 brains) and in *dTBCE* KK + *Mad2* RNAi brains: 0.75 ± 0.11 (*n*=10690, 4 brains). C) Selected live imaging time series in control, dTBCE-depleted, Mad2-depleted or dTBCE+Mad2 depleted dividing Nbs expressing β-tubulin-GFP. The genotypes are indicated on the left. NEBD and anaphase onset are indicated. Scale bar: 10μm. Time is min: s. Dot plot (± s.d.) showing mitosis duration in control (5.1 ± 0.6 min, *n*=67), *dTBCE* KK RNAi Nbs (6.1 ± 0.8 min, *n*=45), Mad2 RNAi Nbs (5.1 ± 0.9 min; *n*=13), and in *dTBCE* KK+ *Mad2* RNAi Nbs (5.3 ± 0.8 min, *n*=30). ***: *P*<0.0001, n.s.=not significant (Wilcoxon test). E) Anaphase neuroblasts stained for α-tubulin (red) and phospho-Histone H3, (PH3, green) to visualize lagging chromatids, indicating defective kinetochore/MT attachments. The genotype is indicated on top of each image. Control (0/21), *dTBCE* KK RNAi (0/17), Mad2 RNAi (0/11) did not show chromosome segregation abnormalities. In fixed preparations, 30% of the cells (4/17) depleted for both *dTBCE* KK and *Mad2*, exhibited lagging DNA. Scale bars: 10μm

**Figure S3. dTBCE is enriched before tubulin in the nuclear space** A) Selected frames of mitosis in a flattened *Drosophila* larval Nb expressing dTBCE-GFP and mCherry-α-tubulin. dTBCE-GFP starts to be enriched in the nuclear space at NEBD and decorates the mitotic spindle region during metaphase. B) Selected frame of a cell entering mitosis in a flattened *Drosophila* larval Nb expressing dTBCE-GFP and mCherry-α-tubulin after treatment with 20 μM colchicine to fully depolymerize the MT cytoskeleton. dTBCE-GFP maximum enrichment (time 00:30) in the nuclear space appears before tubulin enrichment (2:00). Time is min:s. Scale bar: 10 μm.

**Video 1. Embryonic localization of dTBCE-GFP expressed under the control of a poly ubiquitin promoter in a *dtbce* genetic background**

Microtubules (RFP-α-tubulin) are displayed in red and dTBCE-GFP is displayed in green. Scale bar: 10 μm. Time is min: s.

**Video 2. Localization of overexpressed dTBCE-GFP in an early dividing embryo**

Microtubules (RFP-α-tubulin) are displayed in red and overexpressed dTBCE-GFP is displayed in green. Scale bar: 10 μm. Time is min: s.

**Video 3. Localization of overexpressed dTBCE^-mutCG^GFP in an early dividing embryo**

Microtubules (RFP-α-tubulin) are displayed in red and overexpressed dTBCE**^mutCG^**-GFP, harboring a mutated CAP-Gly motif, is displayed in green. Scale bar: 10 μm. Time is min: s.

**Video 4. Localization of overexpressed dTBCE^mut^NLS in an early dividing embryo** Overexpressed dTBCE**^mutNLS^**-GFP, harboring a mutation of the putative NLS, is displayed in monochrome. Scale bar: 10 μm. Time is min: s.

**Video 5. Localization of overexpressed dTBCE^HMF-^GFP in an early dividing embryo** Overexpressed dTBCE**^HMF^**-GFP, harboring a deletion of 4 amino acids downstream of the CAP-Gly motif (that is found in human HMF patients), is displayed in monochrome. Scale bar: 10 μm. Time is min: s.

**Video 6. Localization of EB1-GFP in a control mitotic Nb** Scale bar: 10 μm. Time is s

**Video 7. Localization of EB1-GFP in a dTBCE knock down mitotic Nb** Scale bar: 10 μm. Time is s.

