## Supplementary figures and images for "*Drosophila* dTBCE recruits tubulin around chromatin to promote mitotic spindle assembly"

### Supplementary Figure S1

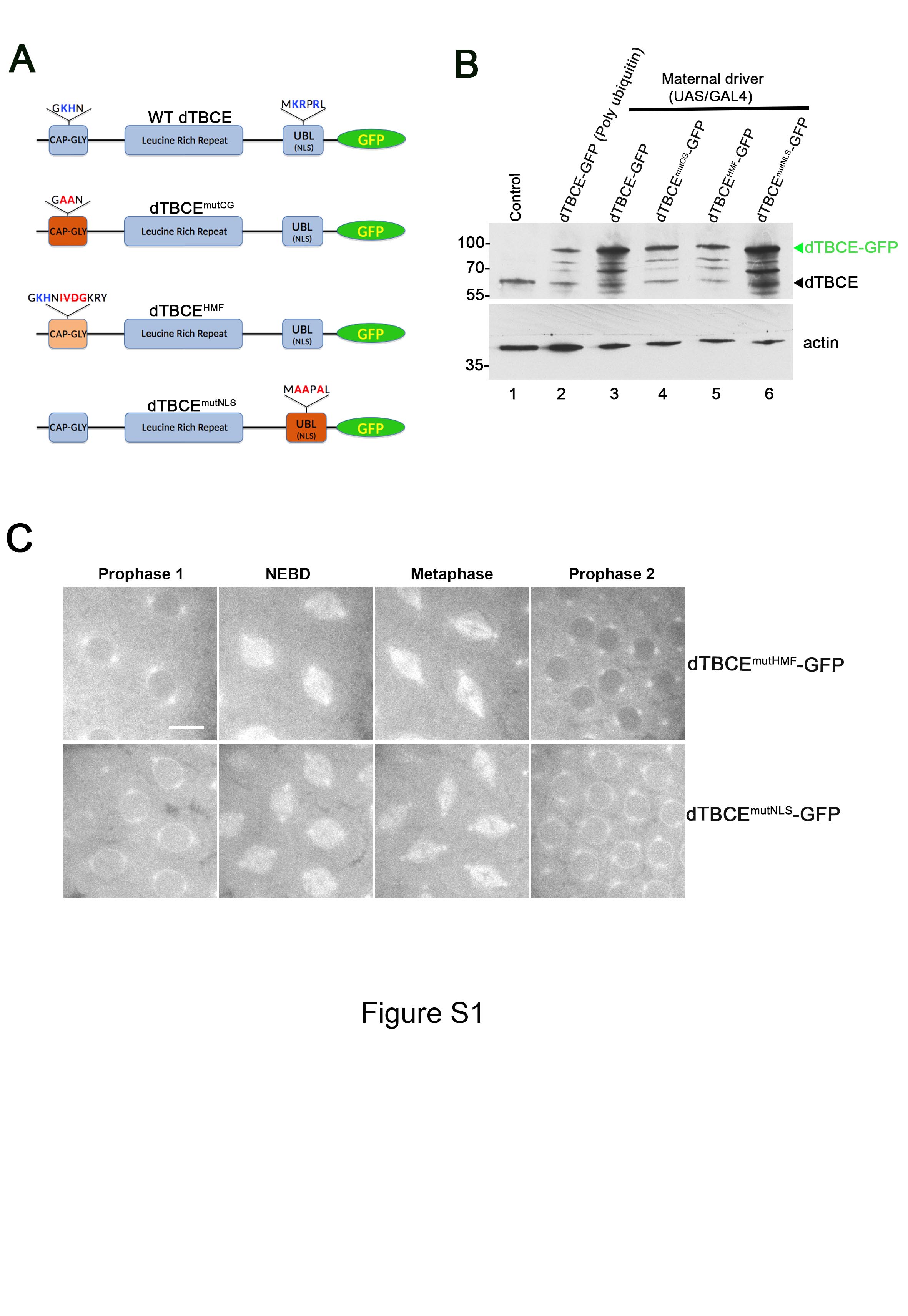

### Supplementary Figure S2

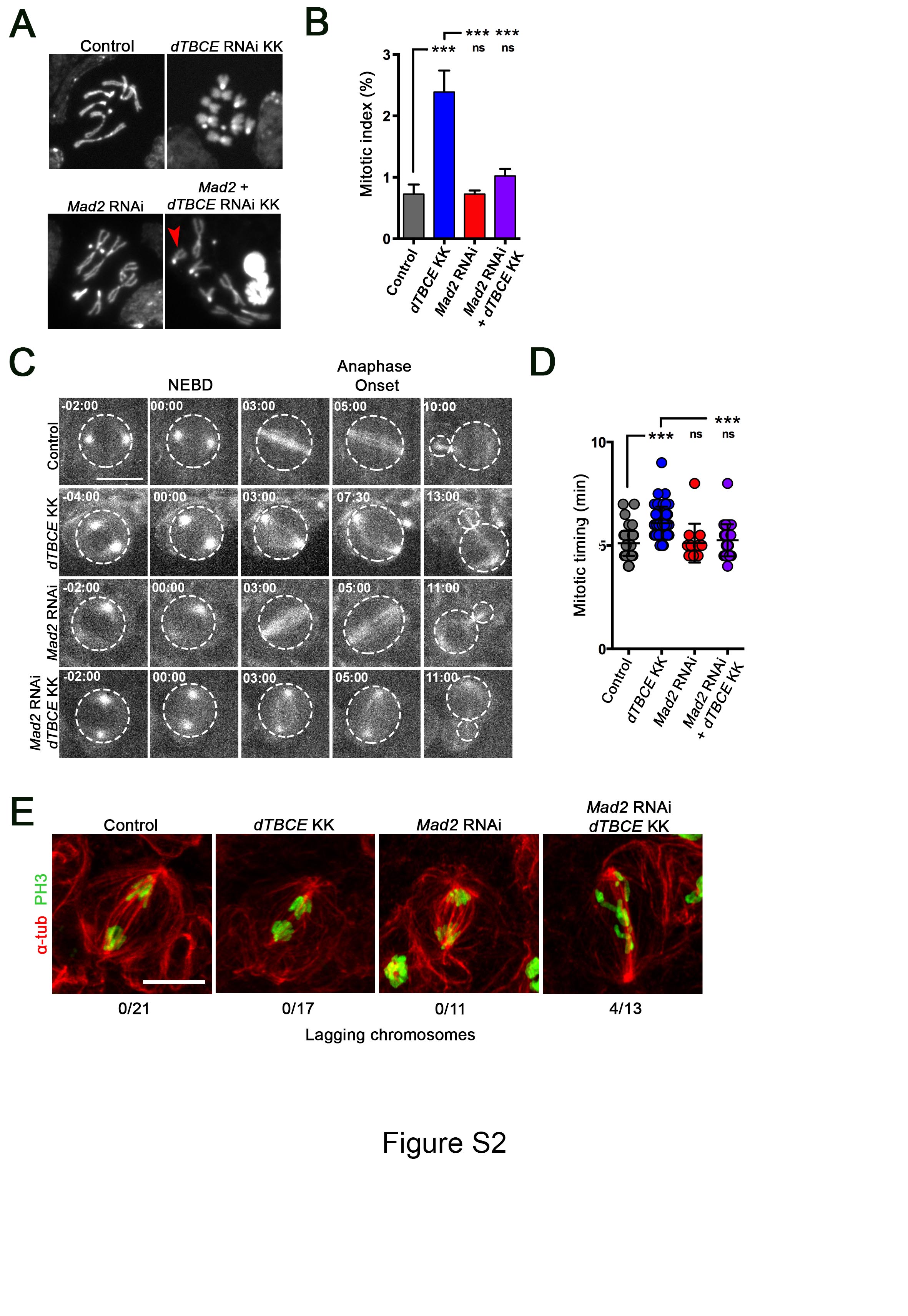

### Supplementary Figure S3

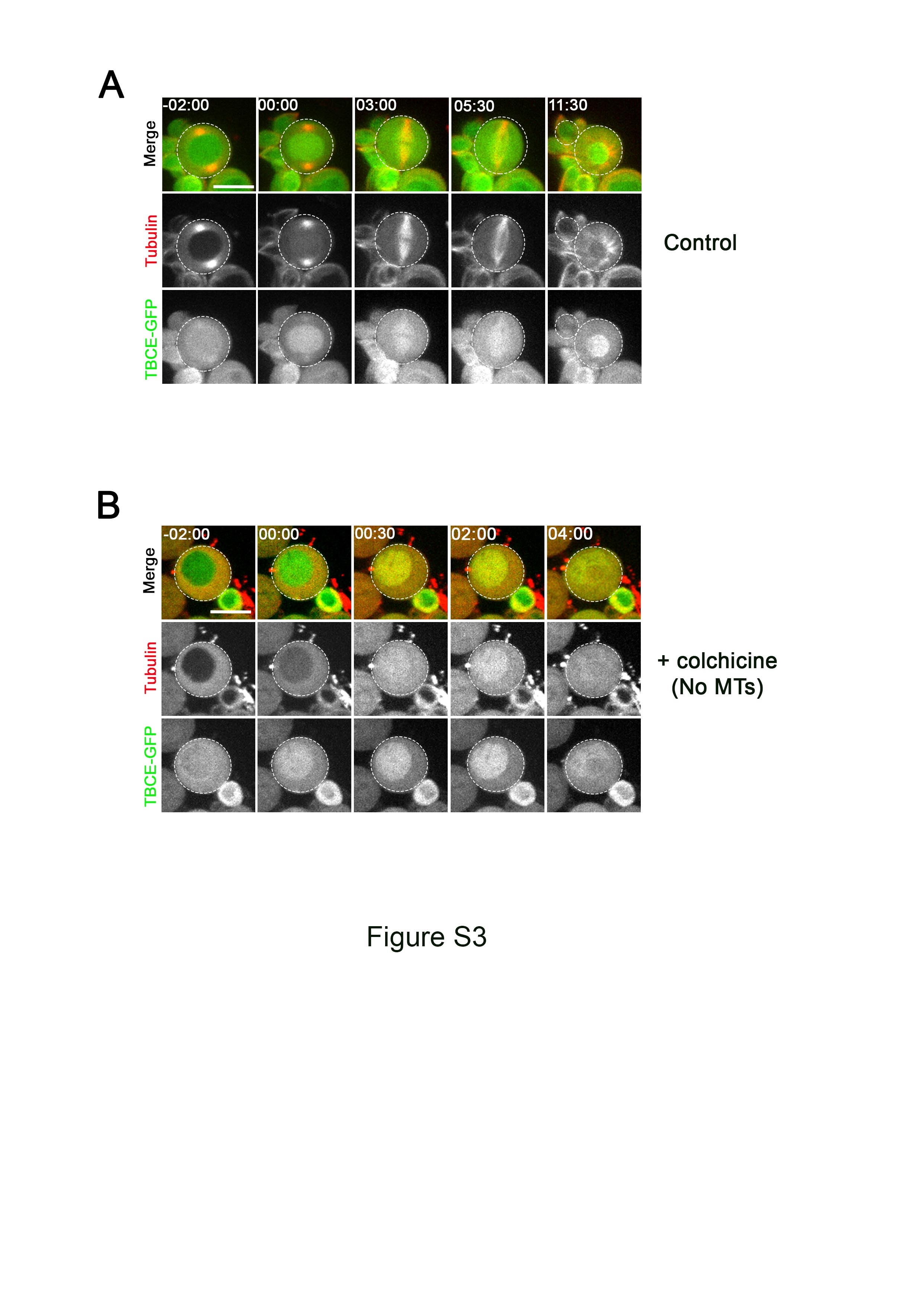
